# Machine learning, molecular docking and simulation studies reveal Lomitapide, Lodipamide, Zafirlukast, Netupitant and Salmon Calcitonin as potent drug molecules against Chagas disease

**DOI:** 10.1101/2024.08.06.606752

**Authors:** Kavya Singh, Navjeet Kaur, Ashish Prabhu

**Affiliations:** Banasthali Vidyapith, Banasthali-304022 (Rajasthan), India; Department of Surgery, University of Oklahoma Health Sciences Center, Oklahoma City, Oklahoma-73104, USA; Department of Chemistry & Division of Research and Development, Lovely Professional University, Phagwara-144411 (Punjab), India; Department of Biotechnology, NIT Warangal, Warangal, India

**Keywords:** Cruzain protein, Docking, Molecular Dynamic Simulation, Machine Learning

## Abstract

*Trypanosoma cruzi* (*T*. *cruzi*), a protozoan parasite, is the pathogen that causes the hazardous disease Chagas, sometimes referred to as American trypanosomiasis. Currently, there are just two drugs commercially available to treat this fatal condition. So, there is an urgent requirement to create novel, safer and efficient anti-Tc drugs. In this study, we introduce a robust experimental strategy that integrates machine learning with molecular docking and simulation studies to identify the most potential drug candidates from the DrugBank dataset for treating the Chagas disease. In a machine learning method, different classifiers (Naïve Bayes, Random Forest, SMO, and C4.5) were used to train the model on the PubChem dataset and the most effective model (C4.5 algorithm) was then chosen and tested on the DrugBank dataset (containing FDA-approved and investigational drugs). The C4.5 algorithm-based machine learning model with an accuracy of 65% predicts the possible drug candidates (a total of 280 drugs were predicted). AutoDock4.2 software was then used to dock the predicted compounds that had a confidence interval of 80% and above. As a result, 47 predicted drugs were docked, and the best of the 5 drugs based on their docking score were chosen for performing the MD simulation studies. MD simulations (100 ns) were conducted for each of the protein-ligand complexes, which produces an average RMSD score (2.23 Å). The RMSF value of the cruzain (Cys**_25_**) binding pocket, after interaction with the five compounds were found to be less than 3.0 Å. Therefore, based on the molecular docking and simulation studies, we found out that Lomitapide, Lodipamide, Zafirlukast, Netupitant and Salmon Calcitonin have shown a stronger binding affinity with the crystal structure of cruzain (Cyst_25_) molecule, making them potential effective inhibitors. Hence, we could anticipate that our five computationally validated drugs will soon be available at the clinical trial stage and eventually be made accessible to the necessary Chagas disease patients.

## 1. Introduction

*Trypanosoma cruzi* (*T. cruzi*), a protozoan parasite, is the pathogen that causes Chagas, often referred to as American trypanosomiasis (Prata, 2001). Triatominae insects or “Kissing bugs” are the main vectors of transmission. Symptoms caused by this disease change over a period of time. Symptoms are usually either absent or are minor and may include headache, high temperature or local swelling at the bite site. *T. cruzi* has many modes of transmission: consuming food tainted with *T. cruzi*, for instance making contact with urine or faeces of infected parasites (Andrade, et al., 2014; Coura and Borges-Pereira, 2010). About 30% of people with persistent infections experience cardiac changes, and up to 10% of people experience changes related to the nervous system, the digestive system, or both. In some later stages, this fatal disease can lead to sudden deaths due to progressive heart failure or cardiac arrhythmias, which are brought on by the destruction of the cardiac muscle and neurological system (Andrade, et al., 2014; Marin-Neto, et al., 2007).

It has been estimated that around 6 million to 7 million people are affected by this *Trypanosoma cruzi*, the parasite that is responsible for causing the Chagas disease (Lidani, et al., 2019). This fatal disease can be prevented through a number of methods. Numerous experimental therapies have shown potential in animal models. These include blockers of squalene synthase, blockers of cysteine protease, dermaseptins extracted from frogs, the sesquiterpene lactone dehydroleucodine (DhL), that have an impact on the development of cultivated epimastigote-phase *Trypanosoma cruzi*, blockers of purine absorption, and blockers of enzymes associated with trypanothione metabolism. Megazol drug seems to be more effective than benznidazole in treating Chagas disease, however, it has not been tested in human beings. Hopefully, after the *T. cruzi* genome is sequenced, new therapeutic targets will become apparent. Chagas vaccine (TcVac3) has been only studied in mice (Barr, 2009; Carod-Artal, 2013; Rassi, et al., 2012).

It is not only present in Latin America, but it is increasingly spreading among different countries such as Europe, Japan, Australia, and North America mainly due to migration. This parasite *T. cruzi* is responsible for causing 20,000 annual deaths, infecting more than 10 million people globally. The treatment cost for curing this particular condition stays significant. In Colombia (2008), it was projected that the yearly expense of medical care for all patients with this condition was around US$ 267 million. Application of insecticide for controlling vectors would cost an annual expenditure of around US$ 5 million –less than 2% of the healthcare expenses (Coura and Borges-Pereira, 2010).

This disease has been prevailing for the last few decades. It was initially spread across Latin America and caused a very low life expectancy rate. Today with the help of several advanced techniques, life expectancy has increased dramatically and also several cures are available for this disease (Barr, 2009). In this new era, people have found several techniques to fight against the disease and also, they have overcome the attached social issues with the disease.

In our research, we are applying ML to predict several new possible drugs and to validate the drugs predicted, we are using Autodock 4.2 and GROMACS software. Using machine learning we are training our model with the inhibitors of the disease caused by *T. cruzi* (PubChem dataset). The DrugBank containing both approved and investigational drugs are then used as a test dataset by the trained model for making predictions. The newly predicted drugs are then validated using the Autodock 4.2 software to make sure that these drugs bind to the same protein active site as the previously accepted drugs does. Furthermore, MD simulations (100 ns) were conducted for each of the protein-ligand complexes, that produces an average RMSD score. The RMSF score of cruzain (Cys**_25_**) binding pocket, upon interaction with the lead molecules was also calculated. Thus, based on the molecular docking and simulation studies, we found out that Lomitapide, Lodipamide, Zafirlukast, Netupitant and Salmon Calcitonin have shown greater binding affinity with the crystal structure of cruzain (Cyst_25_) molecule, making them potential effective inhibitors. Hence, clinical trial studies can be conducted to further analyze these five obtained lead compounds, which can be used in the treatment of Chagas disease.

## 2. MATERIAL AND METHODS

### 2.1. Dataset

PubChem dataset contain around 10822 chemical molecules as two activity sets and they are the inhibitors of *T. cruzi*. These datasets can be found in the bioassay part of the PubChem repository of NCBI, and they possess the AID number as AID 651739 and AID 651740 for identification purposes (Wang, et al., 2012). The molecules are divided into three distinct groups as actives, inactives, and inconclusive.

DrugBank database containing more than 4900 drugs is divided into numerous categories, such as drugs in trial phases, drugs that have been approved, and drugs that have been withdrawn. In this database, over 45% of the drugs have been approved for the usage in a variety of medical conditions (Wishart, et al., 2008). The approved and investigational drugs which is around 2388 have been the subject of our study. These drugs were obtained in the SDF format and, following several processing steps, the resultant description was served as a test model for the training model. The training model was built using the database containing the *T. cruzi* inhibitors. Subsequently, some of the potential drugs have been predicted by the model.

### 2.2. Processing Dataset

These datasets are in the SDF formats. So, to train the model, we need to generate the attributes present in the SDFs. In this case, we have used the PowerMV Description Generator Software. Here, the data obtained in the SDF format is converted into CSV files, which are then utilized to create the training and test datasets for developing the machine learning models (Liu, et al., 2005). These CSV files having both actives and inactives are divided into 80% containing training datasets and 20% containing test datasets. This whole process of splitting is arbitrary. Self-written Python code is used to split the data as per the requirements.

The following FASTA sequence of the cruzain was modeled using the SWISS– MODEL (Biasini, et al., 2014; Guex and Peitsch, 1997; Kiefer, et al., 2009; Schwede, et al., 2003). This provided the predicted structures for the cruzain based on its FASTA sequence. Then, with reference to the QMEAN Score4, we took the best possible prediction among all the predicted structures and the docking of the known as well as the predicted drugs was done based on this structure (Sayers, et al., 2021).

### 2.3. Machine Learning

Machine learning is a part of artificial intelligence that allows us to predict some important features of the datasets after training the model (Mitchell, et al., 1990). Using machine learning, we can think of both classification and regression. It is dependent only upon the algorithm, we are implementing to it and the presentation of the datasets. Here, in this case, we have implemented the classification algorithms.

#### 2.3.1. Classification Algorithm

Classification is a sort of supervised learning which allows a computer system to pick up the knowledge from a dataset that includes specific and useful information. The algorithmic process of classification involves assigning an input value based on the dataset’s descriptions (Liu and Wu, 2012). Therefore, to do this, a mathematical classifier that can label the instances described by the attributes with the appropriate class (Actives and Inactives) is required. During this procedure, the training model is trained on a dataset with pre-assigned classifications and this allows the model to be able to run on distinct datasets so as to classify them based on the current instances (Wang, et al., 2014). In machine learning, there are several algorithms used for classification purposes. In our research, we have compared the outcomes from the classifiers that are Naïve Bayes, Random Forest, SMO, and C4.5.

#### 2.3.2. Training Model

80% of the initial data set is used to prepare the training model. The dataset has been entirely classified, enabling the computer to learn and identify the associations among several attributes. The model is trained using the algorithm coupled with cross-validation. In this instance, suppose the cross-validation is n set with n-folds, then it will split the training data into n parts, of those parts, n-1 will be utilized as a training dataset, and the other one will be utilized for validating the remaining dataset. This procedure of iteration continues for n iteration period of time. In this research, we have utilized a 10-fold cross-validation approach and it is selected based on the dataset’s size (Browne, 2000).

Here, we have two approaches to calculate the misclassification expense with the imbalanced data. The first approach is to define the algorithm as cost-sensitive and then begin with the rest of the settings (Sen and Getoor, 2008). Another method is the usage of a wrapper that helps to transform the basic classifiers into more cost-sensitive ones. This second one is mainly known as Meta-Learning. In this case, at first, using bootstrap aggregating on the decision trees, it estimates the reliable probability. On the basis of that, it relabels the attributes in the training model. Then, these are used in building a cost-insensitive classifier. This also helps in avoiding the algorithm to overfit the dataset and even it reduces the variances of the dataset. Among all the classifiers, we have taken the use of meta-learning in only C4.5 because for two main reasons: one is that it tends to get overfit with the datasets sometimes, and the other is that the meta cost works best with the unprimed trees (Domingos, 1999).

In the case of naïve Bayes, Random Forest, and SMO, we have used the cost-sensitive classifier which utilizes the cost-insensitive procedure for the prediction of the probability estimates of the test cases, and then by utilizing this it predicts the class labels for the test data. In our study, we have divided the datasets into two distinct classes, i.e. actives and Inactives. Thus, we choose the 2X2 matrix, which is frequently utilized in binary classification.

#### 2.3.3. Independent Validation

There are several ways to validate the binary classifiers. True positive rate (TPR) is calculated as the proportion ofthe actual actives to that of the predicted positives (TP/TP+FN), whereas false positive rate (FPR) is computed as the proportion of the predicted false actives to that of the actual inactives (FP/TN+FP). The model’s performance with respect to the actual values can be measured by accuracy, which can be computed as (TN+TP/TN+TP+FP+FN). Sensitivity, which is calculated as (TP/FN+TP), shows how well the model can detect positive outcomes, while specificity, calculated as (TN/TN+FP), demonstrates the model’s capacity to detect the negative outcomes (Jamal and Scaria, 2013).

A model with a low rate of errors has a high level of specificity and sensitivity. Balanced classification rate (BCR) computed as 0.5*(specificity+sensitivity), is the average of the sensitivity as well as specificity that gives the accuracy of the model used on an unbalanced dataset. Mathews correlation coefficient (MCC), whose range of values varies between −1 to 1, is another tool used in addition to the BCR. The representation of the FPR to TPR ratio is shown visually by the receiver’s operating characteristic (ROC) curve. In this instance, the FPR and TPR are positioned on the x-axis and y-axis, respectively. The area that lies under the curve displays the classifier’s probability prediction and its capacity to classify the arbitrarily selected instance into the appropriate one (Periwal, et al., 2012).

### 2.4. Docking of the Predicted Drugs

Our ML model predicted over 280 drugs that may be useful in treating Chagas disease. Two available drugs i.e., benzimidazole and nifurtimox for this disease were predicted accurately by our model with a confidence of 80% (Bermudez, et al., 2016).

Using the AutoDock4.2 software, the compounds that were predicted with more than 80% confidence were docked. So, we docked 47 drugs, and from the best of it, 5 drugs were taken into consideration which have good binding energy and stability with the target molecule cruzain. Autodock 4.2 is the docking software and also it is one of the most cited software in this area. It is most effective for the study of protein interactions with other compounds. This software is maintained mainly by the Scripps Research Institute (Huey and Morris, 2008; Morris, et al., 2009; Trott and Olson, 2010).

Then, after the docking, the most important is the visualization. The visualization is carried out using the PyMOL software. In this case, we have looked upon the pictorial representation of each docked compounds (Delano, 2002).

### 2.5. MD simulations studies

Based on their binding energy in complex formation with the cruzain protein, the top most ligands from the docking studies as well as from the machine learning experiments were subjected to molecular dynamics (MD) simulations.

The AMBER 14 software program was used to perform the simulations. The molecular structures of the proteins as well as ligands were reconstructed by utilizing the generic AMBER force field 99SB and AMBER force field (GAFF), respectively. To neutralize the ligand-protein interactions, the necessary counter ions were utilized. Following that, the products are being solvated using a TIP3P water box of 13 mL. The parameters (“top and crd”) were saved for performing the simulations. Atomic collisions were avoided by minimizing the complexes through the use of steepest decline and conjugate gradient approach for 2500 and 10,000 steps, respectively.

The systems were gradually heated until they exceeded 300 K, and then relaxed during the equilibration step before being put into service. The MD simulations were performed in 100 ns. In order to examine trajectories, the CPPTRAJ module of AMBER 14 was employed.

### 2.6. Binding free energy/MM/GBSA calculations

To reliably assess the ligand-protein interaction relationship, the MM/GBSA technique is performed. The intricate system’s free binding energies were calculated by using the prime module (Jacobson, et al., 2004). The MM/GBSB method (Molecular Mechanics, Generalized Born Model, and Solvent Accessibility) was used to compute the ligand binding energies and strain energies of the docked compound with cruzain protein. Polar solvation energies, non-polar solvation energies, and prospective energies make up the free-binding energy (Bhardwaj, et al., 2021). The OPLS-2005 force field, as well as the SGB solvation equations for polar solvation (GSGB), non-polar solvation (GNP), and Molecular Mechanics Energies (EMM), collaborate with the MM-GBSA, which gathers a variety of nonpolar solvent accessible surface area and van der Waals interactions. Binding energies are calculated from the succeeding equations.

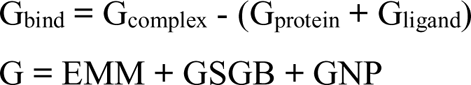

Where, G_complex_ = complex/system energy

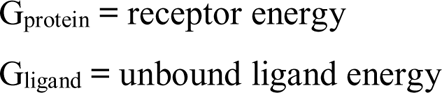

Molecular mechanics energies are reflected by EMM, the SGB solvation term is indicated by GSGB, and the nonpolar solvation model is indicated by GNP (Lokhande, et al., 2021).

## 3. RESULTS AND DISCUSSIONS

### 3.1. Machine learning

In this research, as mentioned, the models were trained using the Classifying algorithm, and the most effective model was then used to test and predict drugs using the DrugBank dataset. Here, we haven’t utilized the unsupervised learning for separating out the datasets because doing so it might significantly weaken the dataset rather than creating it better.

Among the four algorithms, C4.5 was the best on the dataset. It has demonstrated an accuracy of around 64.96% (Table 1). Against this model, we have utilized the drugs found within the DrugBank database to make predictions for the identification of possible drugs that may be used to treat the Chagas disease. Then following this algorithm, the SMO algorithm comes with 63.72% accuracy when tested against the test set which is 20% of the full dataset. After the SMO, the algorithm Random Forest and Naïve Bayes follow this testing pattern with the accuracy of 60.48% and 58.17%, respectively. The C4.5 algorithm has presented around 453 False Positives and 733 True Positives when tested with the test set of 2164 compounds. The C4.5 also has the highest MCC with 0.282 and maximum BCR i.e., 64.63%. Based on the independent validation, the algorithm that produces the lower False Positives and the greater number of True Positives is said to be the most successful model for the drug prediction purposes **(Figure 2)**. When these outcomes are compared to the remaining portion of the dataset, it was found that C4.5 algorithm has performed far better than the other algorithms as it has met both of the above criteria.

**Figure 1:**
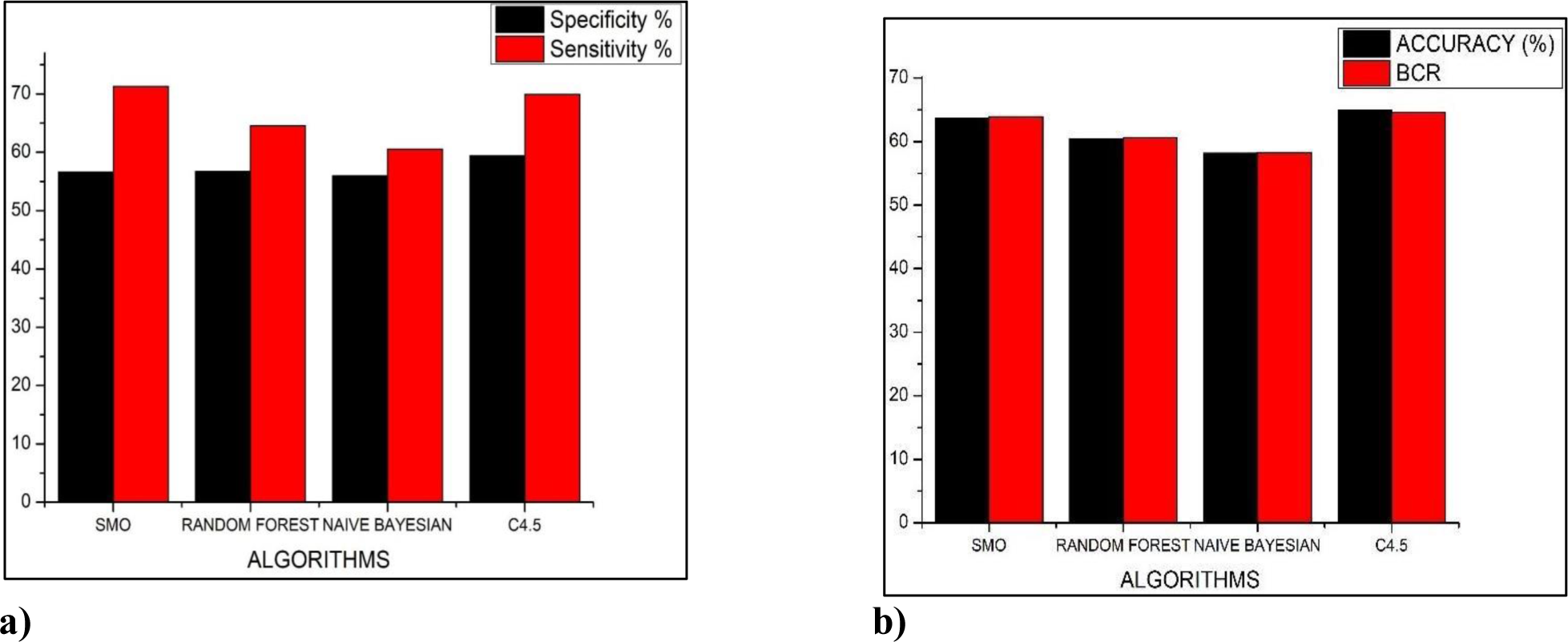
(a) Shows the Comparison of the Algorithm’s Specificity and Sensitivity used for the Preparation of the ML Model. (b) Shows the Comparison of the Algorithm’s Accuracy and BCR used for the Preparation of ML Models.

**Figure 2:**
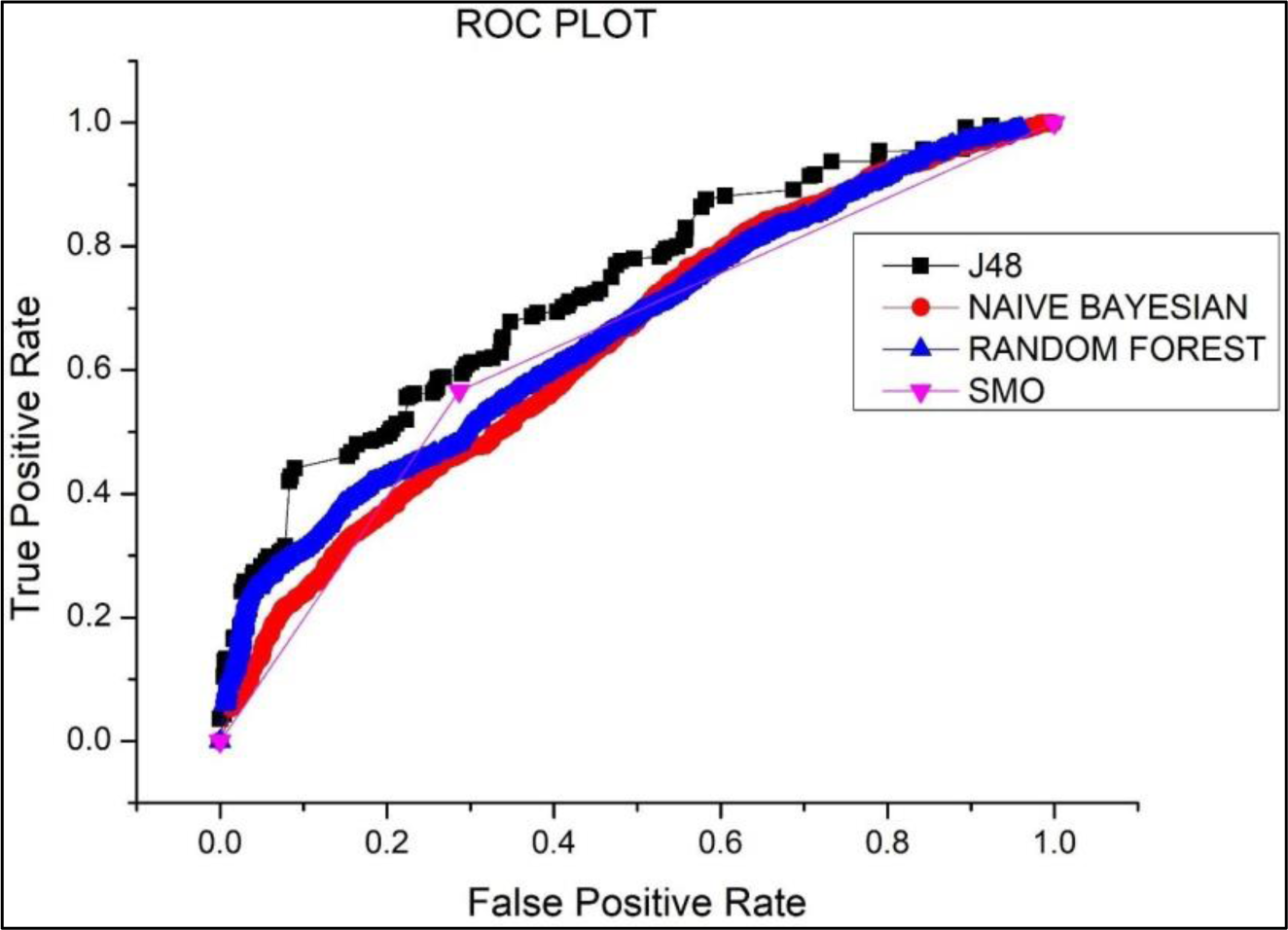
Shows the ROC AUC curve of different algorithms using the 10-fold cross validation.

**Table 1.** – Shows the comparison of the effectiveness of the applied Algorithms for the preparation of the machine learning model.

| ALGORITHMS | SPECIFICITY | SENSITIVITY | ACCURACY | BCR | MCC | ROC |
| --- | --- | --- | --- | --- | --- | --- |
| SMO | 56.59% | 71.3% | 63.72% | 63.94 | 0.282 | 0.639 |
| RANDOM FOREST | 56.68% | 64.53% | 60.48% | 60.6 | 0.213 | 0.663 |
| NAÏVE BAYESIAN | 55.96% | 60.53% | 58.17% | 58.24 | 0.165 | 0.642 |
| C4.5 | 59.37% | 69.9% | 64.96% | 64.63 | 0.294 | 0.697 |

**Table 2:**
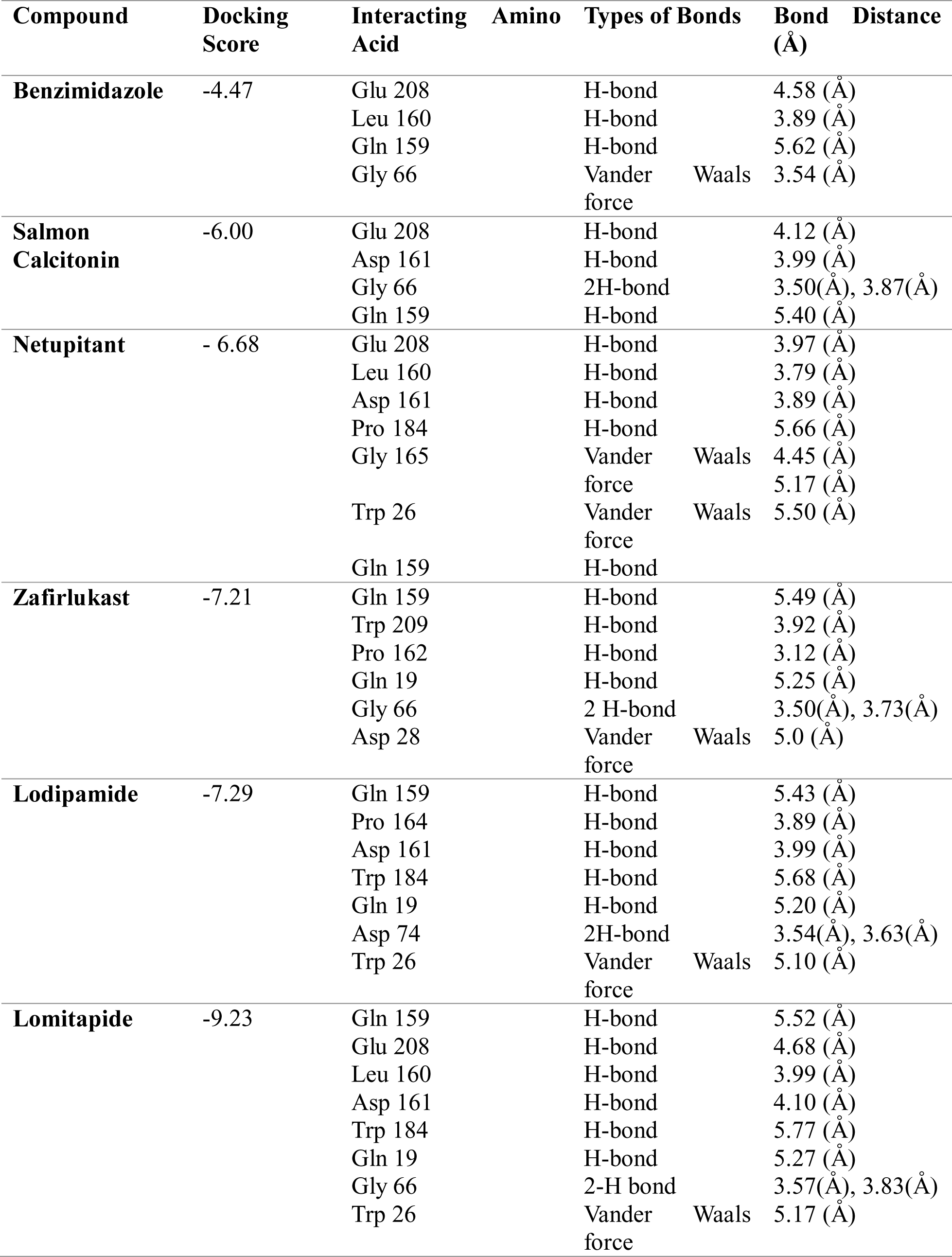
Drug compound docking analysis with cruzain molecule.

The model built using the C4.5 algorithm predicted about 280 drugs out of 2388 drugs that were taken from the DrugBank database. All of the drugs that have been under investigation and/or are currently approved for the treatment of Chagas disease are included in this list of 280 drugs.

Here in this research, AutoDock4.2 software was used to dock the predicted compounds having confidence interval of 80% and above. As a result, we docked 47 drugs, and the best of the 5 compounds were chosen. The docking results have been kept in mind because of their binding energy and their efficacy to interact with the target molecule cruzain.

### 3.2. Cruzain molecule

According to the findings of the conserved domain analysis, cruzain is a member of the cysteine cathepsin proteases family and is mainly responsible for proteolytic activity. It consists of mainly two domains. The first is a-helical (L domain), and the second one is made up of many antiparallel b-sheet interactions **(Figure 3)**. There are two catalytic residues (Cys**_25_** and His**_162_**) present within its active site.

**Figure 3:**
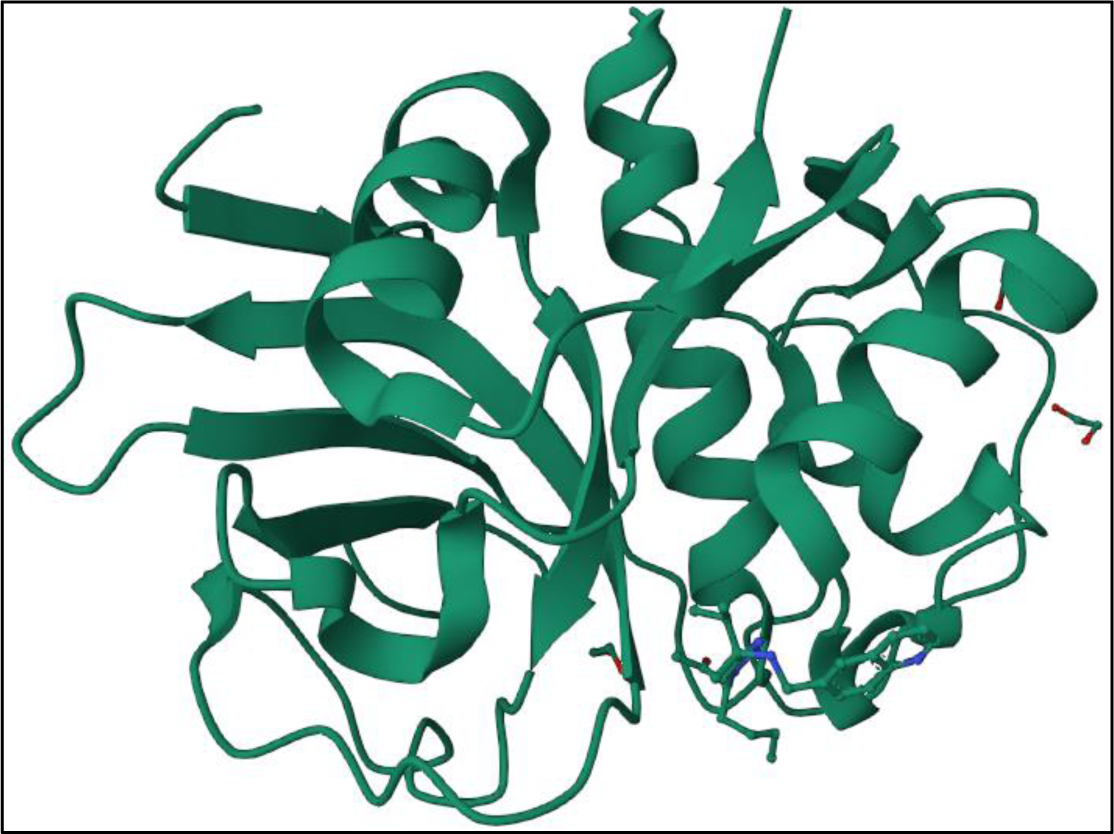
Structure of a cruzain protein.

### 3.3. Free binding energy and compound interaction patterns with cruzain molecule

The cruzain protein was molecular docked for 47 drugs using Autodock 4.2 software. At first, we docked the approved drug i.e., benzimidazole with the predicted structure of cruzain protein. The cruzain protein when docked with the benzimidazole drug (DrugBank ID: DB11989) shows the following binding energy: −4.47eV, which reveals that it had several steric clashes with some of the neighboring strands and therefore, highlighting its ability to prevent the Chagas disease. Overall, the docking analysis suggests that the inhibitor effectively hinders the binding of strands, thus inhibiting the disease.

In account of the docking of the current drugs, we have successfully docked several approved and under investigational drugs predicted by our ML algorithm that have a confidence level of 80% and above. These drugs were taken from the DrugBank database. We discovered that the best drug is Lomitapide (Drug Bank ID – DB08827) using the above reference. The Lomitapide has a binding energy of −9.23 eV **(Figure 4)**, indicating that it has more steric clashes with some of the neighboring strands, suggesting its ability to treat Chagas Disease.

**Figure 4:**
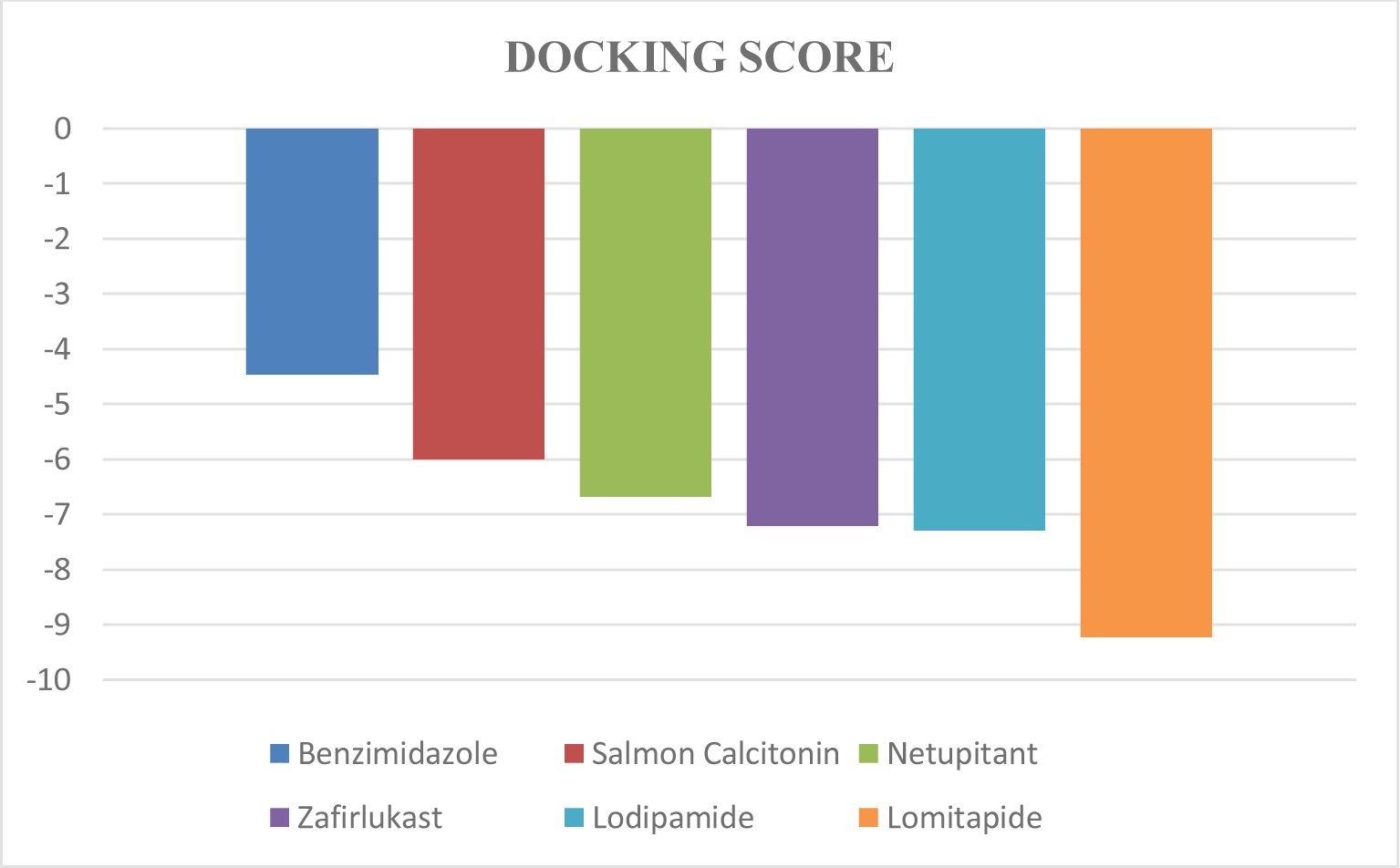
Represents the docking score of the compounds.

In addition to lomitapide (DrugBank ID – DB08827), the second most effective medicine is lodipamide (DrugBank ID-DB04711), which has a binding energy of −7.29. In comparison to the approved drug benzimidazole (DrugBank ID – DB11989), other drugs including zafirlukast (DrugBank ID – DB00549), netupitant (DrugBank ID – DB09048), and salmon calcitonin (DrugBank ID – DB00017) show better potential to control the Chagas disease with binding energy (RMSD) scores of −7.21, −6.68, and −6, respectively **(Figure 4)**.

**Figure 5(a)** depicts the binding location of lomitapide within the curtain protein. With the target protein, lomitapide forms eight hydrogen bonds and one van der Waals force of attraction. As an intermolecular force of attraction, hydrogen bonds keep two or more than two molecules all together. With lomitapide protein, the amino acids Gln 159, Glu 208, Leu 160, Asp 161, Trp 184, Gln 19, and Gly 66 of cruzain molecule form eight hydrogen bonds and one van der waals force of attraction. The Gly 66 forms two hydrogen bonds with lomitapide having a bond distance of 3.53 Å, and 3.83 Å, respectively. Hydrogen bonds generated with lomitapide by amino acids Glu 208, Leu 160, Asp 161, Trp 184, Gln 19, and Gln 159 are present at a distance of 4.68 Å, 3.99 Å, 4.10 Å, 5.77 Å, 5.27 Å, and 5.52 Å, respectively. Trp 26 in the cruzain protein creates a vander walls force of attraction with lomitapide, with a bond length of 5.17 Å. The lomitapide structure is more stable with cruzain protein because of this non-covalent interaction.

**Figure 5:**
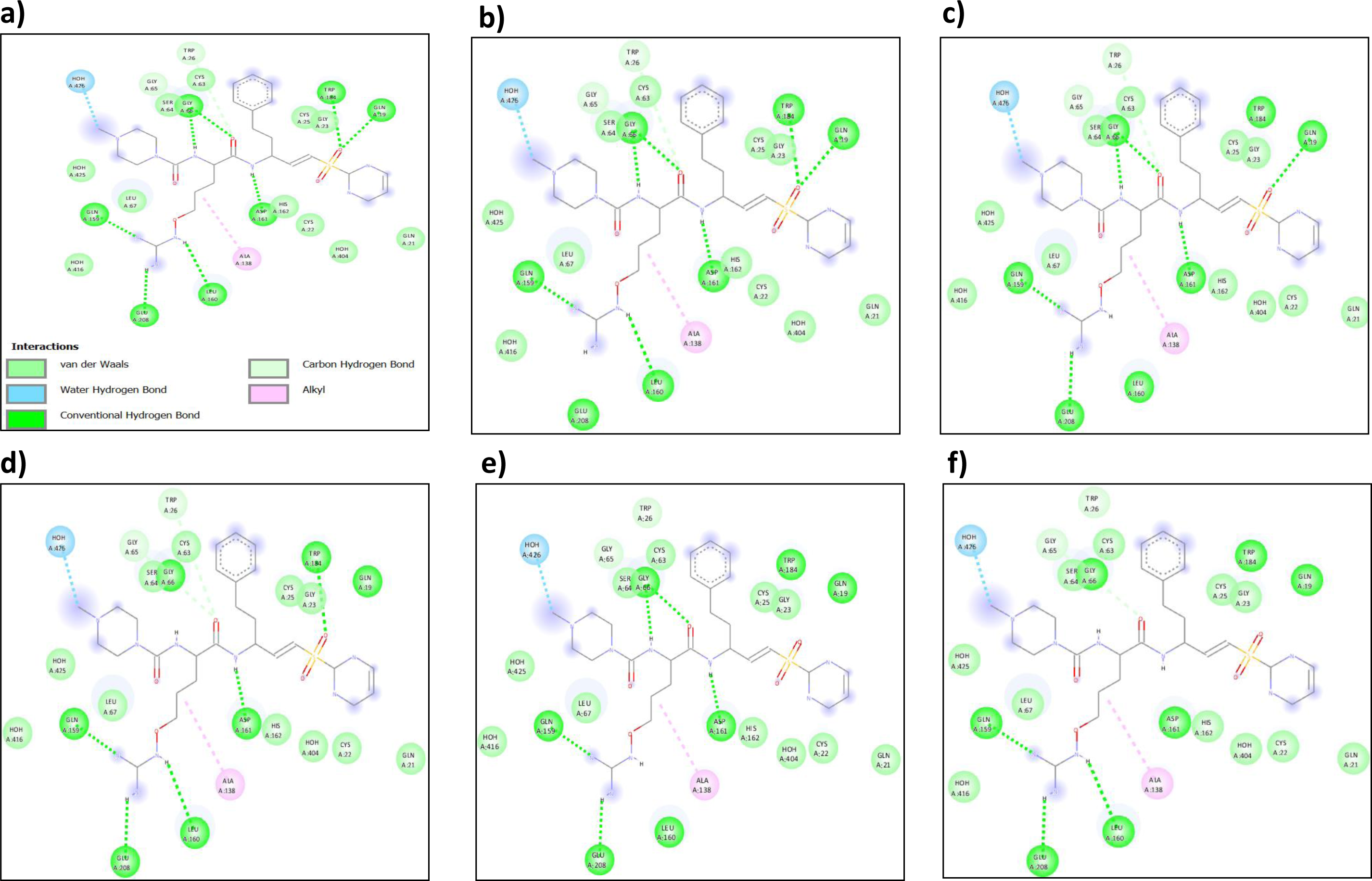
(a) Binding pose of Lomitapide with the cruzain protein binding pocket. (b) Binding pose of Lodipamide with the cruzain protein binding pocket. (c) Binding pose of Zafirlukast with the cruzain protein binding pocket. (d) Binding pose of Netupitant with the cruzain protein binding pocket. (e) Binding pose of Salmon Calcitonin with the cruzain protein binding pocket. (f) Binding pose of Benzimidazole with the cruzain protein binding pocket.

The intermolecular binding interactions between the cruzain structure and another FDA-approved medicine, lodipamide, are depicted in **Figure 5(b)**. Amino acids Gln 159, Pro-164, Asp 161, Trp 184, Gln 19, and Asp 74 forms hydrogen bonds with lodipamide at a distance of 5.43 Å, 3.89 Å, 3.99 Å, 5.68 Å, 5.20 Å, and 3.54 Å, respectively. Lodipamide has a docking score of −7.29 **(Figure 4)**, indicating its ability to treat Chagas disease.

The binding position of zafirlukast within the cruzain structure is indicated in **Figure 5(c)**. With a binding energy of −7.21, zafirlukast produces six hydrogen bonds and one vander wal force of attraction. The amino acid Gly 66 forms two hydrogen bonds with zafirlukast at 3.50 Å and 3.73 Å bond length, respectively. Furthermore, Gln 159, Trp 209, Pro 162, and Gln 19 at a distance of 5.49 Å, 3.92 Å, 3.12 Å, and 5.25 Å, respectively form one hydrogen bond each. At a bond distance of 5.00 Å, Asp 28 is involved in forming vander waals force of attraction with zafirlukast. **Figure 5(d)** illustrates the formation of hydrogen bonds within amino acids Glu 208, Leu 160, Asp 161, Pro184, and Gln 159 present in netupitant at a bond distance of 3.97 Å, 3.79 Å, 3.89 Å, 5.66 Å, and 5.5 Å, respectively. Similarly, residues Gly 165 and Trp 26 **(Figure 5d)** form vander walls force of attraction at a bond length of 4.45 and 5.17, respectively, resulting in the development of a stable complex between the netupitant compound and cruzain molecule. In **Figure 5(e)**, Glu 208, Asp 161, and Gln 159 form one hydrogen bond each with salmon calcitonin molecule with a bond length of 4.12 Å, 3.99 Å, and 5.40 Å, respectively. Moreover, Gly 66 forms two hydrogen bonds at a distance of 3.50 Å and 3.85 Å, respectively. These non-covalent interactions are crucial in determining the stability of a molecule. Salmon calcitonin is a stable molecule that can be utilized to treat people with Chagas disease since it has a binding energy of −6.00 and non-covalent interactions.

### 3.4. RMSD and RMSF calculation

Since the cruzain protein’s rigid structure was used for molecular docking calculations, we implemented MD simulation to examine the stability of the bound state following the lead compound binding inside the cavity of the cruzain molecules. We have simulated the process for up to 100 ns in order to examine the structural stability of complexes made up of protein-ligand. In our study, we have utilized the duration of 100 ns that is considered to be a sufficient time required for the rearrangements of atoms of the cruzain molecule in associations with the ligand molecules.

The Root Mean Square Deviation (RMSD) and Root Mean Square Flactuation (RMSF) scores of Cyst**_25_** were assessed in order to display the thermodynamic structural stability over a period of 100 ns. Furthermore, 1000 trajectories produced by the simulation were recorded and imposed onto the cruzain (Cys25) crystal structure by using the simulation event analysis (SEA), that produces results in ".dat" manner. The RMSD and RMSF graph was created using the compiled data of RMSF and RMSD values. RMSD plot **(Figure 6)** demonstrates that all the reported drugs, namely Lomitapide, Lodipamide, Zafirlukast, Netupitant and Salmon Calcitonin are present within the permissible limit of RMSD with an average value of 2.23 Å, suggests that all the compounds are strongly bounded inside the cavity of the cruzain molecule. Furthermore, the RMSD graph **(Figure 6)** in our study shows that all complex systems equilibrate under the 1 to 3.5 Å RMSD range between 30 and 100 ns, with a mean value of 2.23 Å (RMSD).

**Figure 6:**
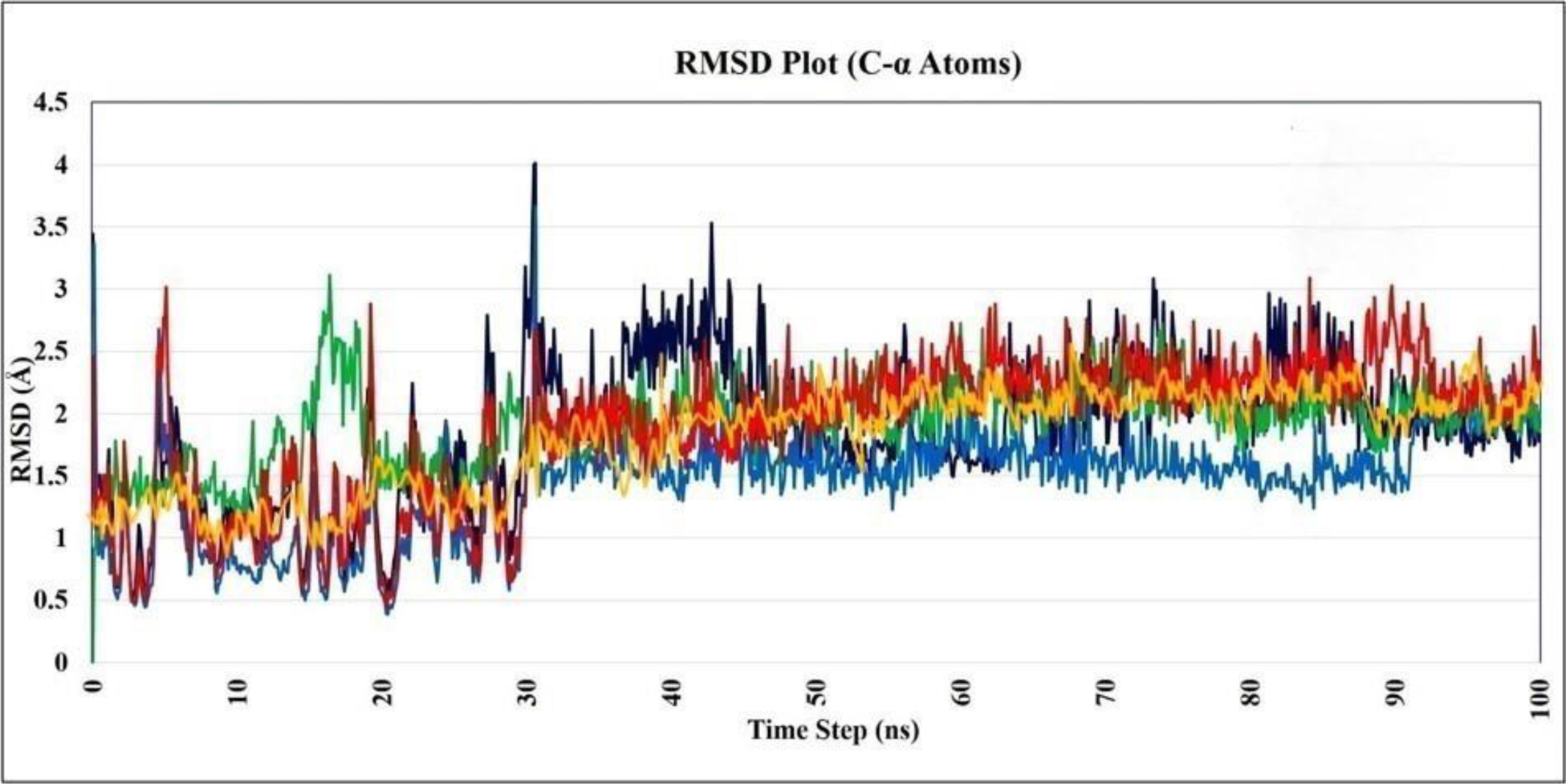
Time dependence of root mean square deviations (RMSDs) of the cruzain (Cys**_25_**) has been depicted after binding with the ligand molecules i.e. Lomitapide (dark blue), Lodipamide (blue), Zafirlukast (green), Netupitant (red) and Salmon Calcitonin (yellow).

On the other side, root mean square fluctuation (RMSF) scores for each residue were calculated in order to assess the binding effectiveness of reported compounds with cruzain (Cys**_25_**) molecule. The mean RMSF value calculated for Cys**_25_** by binding of Lomitapide, Lodipamide, Zafirlukast, Netupitant and Salmon Calcitonin is 1.7 Å since each component varied between the 0.3 Å to 3.2 Å RMSF range **(Figure 7)**, which depicts the minimal variation and maximum stability of cruzain (Cys**_25_**) on binding of lead molecules. Thus, the MD studies have shown that Lomitapide, Lodipamide, Zafirlukast, Netupitant and Salmon Calcitonin exhibits greater potential against cruzain (Cys**_25_**) molecule.

**Figure 7:**
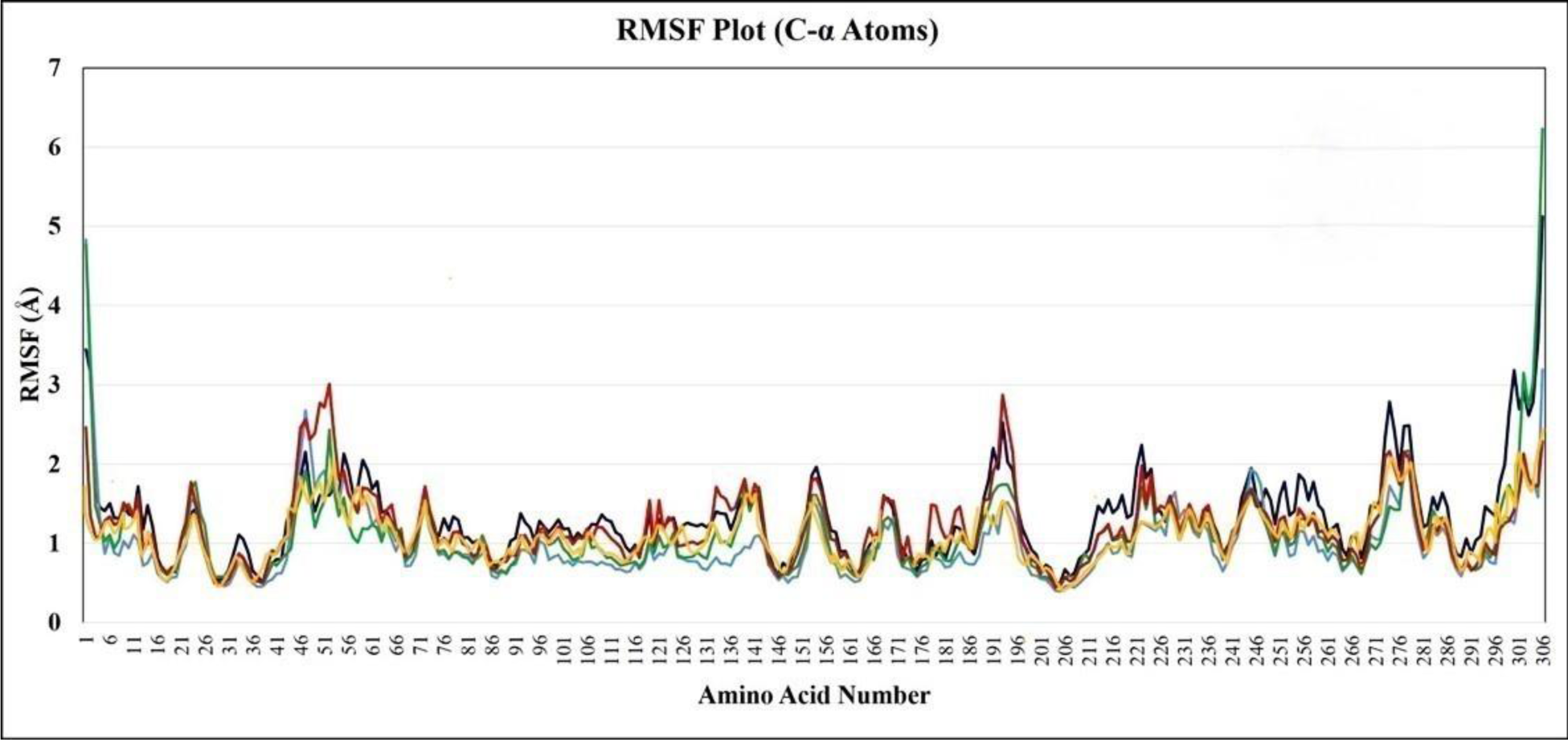
Time dependence of root mean square fluctuation (RMSFs) of the cruzain (Cys**_25_**) has been depicted after binding with the ligand molecules i.e. Lomitapide (dark blue), Lodipamide (blue), Zafirlukast (green), Netupitant (red) and Salmon Calcitonin (yellow).

In addition to this, throughout the duration of MD simulation (100 ns), we have investigated the stability of the binding pocket after the attachment of the ligand to the cruzain (Cys**_25_**) molecule by measuring the RMSF values of each residue that falls within the binding pocket of the Cys**_25_**. The 1000 trajectories produced by MD simulation (100 ns) were recorded and imposed over the crystal structure to measure the RMSF values of each residue present within the binding pocket (Table 3). The RMSF value of cruzain (Cys**_25_**) binding pocket, after interaction to compounds like Lomitapide, Lodipamide, Zafirlukast, Netupitant and Salmon Calcitonin is less than 3.0 Å, which indicates that the binding cavity of cruzain (Cys**_25_**) is fairly stable throughout the course of MD simulation.

**Table 3:** RMSF values of cruzain (Cys_25_) binding pocket residues after binding of ligand molecules.

| **Residues** | **Lomitapide** | **Lodipamide** | **Zafirlukast** | **Netupitant** | **Salmon**<br>**Calcitonin** |
| --- | --- | --- | --- | --- | --- |
| **Gly65\*** | 1.438 | 1.500 | 1.668 | 1.507 | 1.45 |
| **Ser64\*** | 0.848 | 0.847 | 1.199 | 0.900 | 1.88 |
| **Gly66\*** | 0.655 | 0.669 | 0.756 | 0.456 | 2.93 |
| **Cys63** | 0.789 | 0.601 | 0.700 | 0.555 | 1.46 |
| **Trp26\*** | 1.434 | 1.219 | 0.900 | 1.967 | 0.77 |
| **Cys25\*** | 1.387 | 1.674 | 1.204 | 2.224 | 1.87 |
| **Gly23\*** | 1.600 | 1.634 | 2.609 | 2.779 | 2.77 |
| **Trp184\*** | 1.620 | 0.923 | 1.390 | 1.112 | 1.66 |
| **Gln19\*** | 1.465 | 0.939 | 1.307 | 0.567 | 2.99 |
| **Gln21\*** | 1.649 | 1.023 | 1.234 | 1.478 | 1.68 |
| **His162\*** | 1.489 | 1.089 | 1.745 | 1.946 | 1.90 |
| **Asp161\*** | 1.137 | 0.979 | 1.220 | 1.100 | 1.54 |
| **Ala138\*** | 0.800 | 0.659 | 0.723 | 0.959 | 1.88 |
| **Leu160\*** | 1.019 | 0.939 | 0.766 | 0.654 | 1.23 |
| **Glu208\*** | 0.919 | 0.689 | 0.967 | 1.166 | 0.98 |
| **Gln159\*** | 0.785 | 0.590 | 0.298 | 1.765 | 0.55 |

### 3.5. MM/GBSA energies calculation for ligands complexed with cruzain (Cys25)

MM/GBSA has been found to be the most appropriate method to measure the free binding energies that describes the outcomes as hydrophobic, VDW, or solvation elements. The cruzain (Cys**_25_**) ligands i.e. Lomitapide, Lodipamide, Zafirlukast, Netupitant and Salmon Calcitonin were screened and then subjected to MM/GBSA technique for a large period of MD simulation. The computed binding free energies of five complexes employing prime MM/GBS methods are listed in Table 4. The protein-ligand interaction is robust and stable when there is a greater value of negative (less binding energy). Based on the findings, it can be inferred that Lomitapide has the highest binding energy (−161.309 kcal/mol), signifying its strong interaction stability. On the other hand, other ligand molecules like Lodipamide, Zafirlukast, Netupitant, and Salmon Calcitonin also exhibit strong binding free energies with cruzain (Cys**_25_**), indicating that all these compounds form a robust complex with the target molecule. The MM/GBSA outputs indicate that the ligand molecules fulfil the MM/GBSA method to form a robust complex with cruzain (Cys**_25_**), and all the values estimated by MM/GBSA are dynamically favorable.

**Table 4:** Shows the docked structure’s binding energies (kcal/mol) over the course of MD simulation (100 ns).

| Compounds complexed with Cys<sub>25</sub> | DG Bind<sup>a</sup> (kcal/mol) | DG Coulomb<sup>b</sup> (kcal/mol) | DG Bind vdW<sup>c</sup> (kcal/mol) | DG Solv GB<sup>d</sup> (kcal/mol) | Complex Energy<sup>e</sup> (kcal/mol) |
| --- | --- | --- | --- | --- | --- |
| Lomitapide | —161.309 ± 7.029 | 81.804 ± 10.078 | 75.685 ± 2.814 | 93.719 ± 12.234 | —15349.07 ± 66.519 |
| Lodipamide | —158.890 ± 10.155 | 89.720 ± 5.802 | 73.900 ± 4.443 | 98.90 ± 7.909 | —15329.31 ± 67.99 |
| Zafirlukast | —154.870 ± 3.651 | 63.423 ± 7.948 | 75.160 ± 1.685 | 75.758 ± 4.668 | —15350.44 ± 44.619 |
| Netupitant | —150.344 ± 3.337 | 50.436 ± 7.928 | 76.472 ± 2.219 | 54.0820 ± 4.938 | —15406.90 ± 62.801 |
| Salmon Calcitonin | —147.333 ± 3.331 | 49.436 ± 7.919 | 76.387 ± 2.109 | 51.0798 ± 4.927 | —15402.90 ± 62.800 |

<sup>a</sup> MM/GBSA binding free energy.
<sup>b</sup> Coulomb Coulomb energy.
<sup>c</sup> Van der Waals energy.
<sup>d</sup> GB Generalized Born electrostatic solvation energy.
<sup>e</sup> Energy of protein-ligand complex.

## 4. Conclusion

The hazardous disease known as “Chagas”, sometimes called “American trypanosomiasis,” is brought on by the protozoan parasite *Trypanosoma cruzi* (*T. cruzi*). Only two drugs (benznidazole (BZN) and nifurtimox (NFX) are currently available in today’s marketplace to treat this deadly disease. Due to the severe adverse effects of these drugs, patients often stop taking them, which leads to ineffective pharmacotherapeutic effect. So, there is an urgent requirement to create a novel, safer and efficient anti-Tc drugs. In our study, we present a strong experimental approach that combines machine learning with molecular docking and simulation studies to find the most promising drug candidates from the DrugBank dataset to treat the Chagas disease. In a machine learning method, different classifiers (Naïve Bayes, Random Forest, SMO, and C4.5) were used to train the model on the PubChem dataset and the most effective model (C4.5 algorithm) was then chosen and tested on the DrugBank dataset (containing FDA-approved and investigational drugs). The C4.5 algorithm-based machine learning model with an accuracy of about 65% predicts the possible drug candidates. A total of 280 drugs, including those that were already available in the market, were predicted. The predicted drugs with a confidence interval of 80% and above were docked using Autodock 4.2 software. As a result, 47 predicted drugs were docked, and the best of the 5 drugs based on their docking score were chosen. Lomitapide, Lodipamide, Zafirlukast, Netupitant and Salmon Calcitonin displayed a docking score of −9.23, −7.29, −7.21, −6.68, and - 6.00, respectively. Hence, we found that Lomitapide, Lodipamide, Zafirlukast, Netupitant and Salmon Calcitonin have shown a stronger binding affinity with the crystal structure of cruzain (Cyst_25_) molecule, making them potential effective inhibitors.

Each of these five protein-ligand complexes underwent molecular dynamic simulations (100 ns), which produced an average RMSD score of 2.23 Å. This shows the ligand molecule’s strong binding ability inside the cavity of the cruzain molecule and thus resulting in conformational stability. The RMSF value of cruzain (Cys**_25_**) binding pocket, after interaction to compounds like Lomitapide, Lodipamide, Zafirlukast, Netupitant and Salmon Calcitonin is less than 3.0 Å, which indicates that the binding cavity of cruzain (Cys**_25_**) is fairly stable over the course of the MD simulation. Hence, in vivo, in vitro, and clinical studies can be carried out to further analyze these five acquired lead compounds that could be used for the treatment of the Chagas disease.

## Acknowledgements

K.S. is thankful to the MHRD fellowship provided during the PhD tenure.

## Author’s contribution

**KS:** Docking studies, Molecular dynamics, Machine learning, Research design, Data collection, Writing manuscript. **NK:** Manuscript editing and Supervision. **AP:** Manuscript editing and Supervision.

## Conflict of interest

All authors contributed to the article and approved the submitted version. There are no conflicts of interests.

## Data availability statement

The corresponding author can provide the docking structures upon request.

## Notes

### Competing Interest Statement

The authors have declared no competing interest.

